# Theoretical principles of multiscale spatiotemporal control of neuronal networks: a complex systems perspective

**DOI:** 10.1101/097618

**Authors:** Nima Dehghani

## Abstract

Success in the fine control of the nervous system depends on a deeper understanding of how neural circuits control behavior. There is, however, a wide gap between the components of neural circuits and behavior. We advance the idea that a suitable approach for narrowing this gap has to be based on a multiscale information-theoretic description of the system. We evaluate the possibility that brain-wide complex neural computations can be dissected into a hierarchy of computational motifs that rely on smaller circuit modules interacting at multiple scales. In doing so, we draw attention to the importance of formalizing the goals of stimulation in terms of neural computations so that the possible implementations are matched in scale to the underlying circuit modules.

## A. Overview

In this theoretical perspective article, we propose the need for a multiscale information theoretic framework on how to better control an adaptive complex system such as the mammalian nervous system. We start with a brief historical overview of neural stimulation from Galvani’s pioneering work to modern opto-electrical methods. Pointing to control obstacles, we portray why an understanding of the information processing levels of neuronal networks is crucial to the proper design of a control paradigm. We then emphasize the importance of scale-independence and examine the shortcomings of network control without it. This interpretation lays out the need for attention to the dynamic nature of computation and the functional robustness in the light of structure variability thus illustrating the multiscale information processing nature of neuronal networks. We advance the idea that, within this framework, better control can be achieved through targeting the aggregate computational output rather than attempting to control the system at its finest scales.

## B. Control obstacles and failures of neural stimulation

In 1780, Luigi Galvani discovered that an electrical spark causes the twitching of a dead frog’s legs. This discovery was pivotal to the birth of bioelectromagnetism and led the to the idea of controlling action and behavior with electricity^15,122^. A century after Galvani’s famous experiment, Gustav Fritsch and Eduard Hitzig showed that, in dogs, *in vivo* motor cortex stimulation causes limb movement^38^. Electrical neuromodulation as a therapeutic tool has since been recognized as a viable and concrete path to control neurological diseases^41^. Recognition of the implication of the Substantia Nigra’s damaged cells in Parkinson’s disease^59^, better understanding of the operational principles of basal ganglia-thalamocortical loops^1^ and the effective reduction of tremor by lesioning the subthalamic nucleus in monkey model of Parkinson’s^10^ provided the path to tackle Parkinson’s. Following these discoveries and a century after Fritsch and Hitzig’s experiments the first deep brain stimulation (DBS) operation was performed in Parkinsonian patients in 1987 and positive clinical effects were reported a few years later^56,64^.

However, despite the technological advancements in electrical stimulation, control of the behavior has been plagued by improper precision and lack of understanding of the network response. In the case of macro-stimulation (such as TMS – transcranial magnetic stimulation, tDCS – transcranial direct stimulation, DBS), much of the effort is focused on improved targeting of a smaller area. The hope is that via fine-tuned targeting, one would eventually achieve the proper level of control. Nonetheless, electrical micro-stimulation has not shown much promise in precise control of the output of circuits. For example, microstimulation of MT (middle temporal visual area) directional-selective neurons shows variable behavioral efficacy across single trials, where this variability has been ascribed to attention gating^84^ or spatial feature-selective gating^86,104^. A major issue with the electrical micro-stimulation is the imprecision in targeting a specific cell-group (excitatory vs inhibitory, or a given group of inhibitory cells) as well as the nebulous temporal control of the stimulation effect. These imprecisions turn neural control to art rather than exact science. Fine tuning DBS for selection of good stimulus parameters is done mostly with trial and error^124^. Additionally, the timescale within which TMS effects may last can vary over orders of magnitude. Moreover, the effects of TMS or tCS (direct or alternating transcranial stimulation) can easily spread to spatial neighbors of the actual target^26^.

The discovery of optogenetics (use of light to control neuron) in cultured mammalian neurons^127^ triggered the next wave of neuromodulatory attempts through temporally precise stimulation of individual excitatory and/or inhibitory neurons with millisecond resolution^14,128^. An ever-growing list of studies relying on optogenetics stimulation have since followed, including those targeting Parkinson’s^55^, aimed at behavioral conditioning^119^ or fear-conditioning^45^, as well as targeting deep structures (brainstem)^65^, or for inducing fast (*γ*) rhythms^19^, modulating sensory processing^107^ and attempts to control network disorder (hippocampal seizure)^57^. From device engineering perspective, optogenetics’ major challenges include geometrical and mechanical design issues, light delivery stability and precision, optimization of light/power efficiency, heat dissipation^43,123,129^. Other issues, specially for applicability in clinical settings, relate to safe opsin molecular engineering, safe opsin delivery and optical stimulation techniques^49^. For these reasons, optogenetics is still not a viable option for clinical purposes^43,49^. However, the main biophysical challenges are photoelectric artifact (Becquerel effect) and photothermal effects^54^. Specifically, simultaneous optical stimulation and electrical recording leads to photo-electric artifacts which could be incorrectly interpreted as the light-induced rhythmic activity^54,129^.

Many of these optogenetics challenges will be resolved with engineering advances. However, the attempts for control through optogenetics have faced other serious challenges related to the underlying information processing of the systems under study. The biggest hurdle is our limited system level understanding of brain functions^49^. The observed variability of the evoked behavior following transient inactivation of the motor cortex in rats and nucleus interface (Nif) in songbirds^90^ has seriously challenged the assumption that optogenetics can effectively overcome nonspecific cell targeting and temporal imprecision. As proposed, ignoring the indirect effects of downstream circuits^90^ and disregarding the interconnectedness of the complex circuitry are among the key reasons that even a temporally-precise cell-specific control of individual elements (i.e. neurons) of the network can not yield precise control of the network function. The macroscopic behavior of the system (such as network balance of excitation/inhibition) is insensitive to the computational state of individual neurons^28^. This insensitivity is not because the functional symmetry of individual elements transcends to the total state^3^, but because interconnectedness renders many details (at fine scale) to be irrelevant at the large-scale behavior of the system^42^. Thus, attempts for precise control of the system at its fine scale is precisely where it will fail. This failure roots in the breakdown of the constructionist view and its emphasis on the independence of behavior at microscopic and macroscopic scales^3^. In what follows, we emphasize on a the need for a paradigm shift to understand and control the system in a multiscale information theoretic framework.

## C. Levels of information processing

Proper success in augmentation/alteration of the nervous system’s behavior roots in the deep understanding of the complexity of this computational system. Here, by complexity we explicitly refer to a) existence of multiple scales of structure/dynamics, b) although the structural elements of coarser scales can be physically reduced to those of the finer scales, the macro-micro dynamical elements are neither fully independent nor completely coherent, and c) the number of possible states of the system (i.e the amount of information – in bits– needed to define the system) is scale-dependent. Early attempts to describe the nervous system at multiple scales, led to the formation of a tri-level hypothesis of information processing. This approach divides the description of the system into its computational, algorithmic, and implementational subsets after David Marr^78,79^ or to semantic, syntactic, and physical categories according to^97^. While this view in neuroscience and philosophy/cognitive science has had a tremendous impact (specifically in the field of visual neuroscience), the missing links across the tri-levels have been the target of criticisms and a source of observed failures in the description of neural computation at the system level and its connection to behavior. Interestingly, even though Marr initiated this classification of levels, his own studies were always focused on only one level at a time. In fact, Marr’s own quest was initially more focused on the fine scale level^74–76^, but later he became skeptical^77^ and switched to the top, i.e. computational, level^80,101^. The main issue in this approach is reliance on the separation of scales, a notion deeply in conflict with the nature of complex systems^7^. The behavior of the systems harboring hierarchical levels of sub-assemblies is defined by the interaction of the sub-assemblies at higher levels not by details in a given sub-assembly^110,111^. Surprisingly, a recent reversal of interest in the implementational level (mainly due to advances in microscopy and computational tools) has led to the revival of the constructionist view. This approach entails that by taking into account the adjacent biophysical details of cell type categories, their placement in micro-circuitry and their afferent and efferent projections, one could achieve proper control/alteration of neural information processing systems.

The recent surge of interest in the fine level details of both structure^16,50^ and simulation^73^ parallels Marr’s initial emphasis on fine implementational details. This approach has drawn criticism from advocates for his later emphasis on the computational level. The underlying assumption is that through a combined study of in vivo physiology and network anatomy^11^, one can build functional connectomics^105^ and decipher the behavior of the system. In small systems lacking the hierarchical architecture of complexity, separation of scales is justifiable and such an approach may lead to a good understanding of the system’s behavior (such as Dorsophila motion detection^117^), or yield relatively good control of simple behaviors (such as optogenetic control of simple motor in C elegans^62^). The opposing view – mainly driven by system neuroscience investigations of the mammalian neo-cortex – argues that the understanding the underlying details will not resolve the issues relevant to the neural computation of interest^17^. This vantage point suggests that a repetition of canonical computation such as linear filtering^87,102^ or divisive normalization^46,112,130^ serves as building blocks of computation and is the cornerstone of information processing^18^ leading to progressively more complex representation in the hierarchy of sensory processing cortical areas^53^. It is suggested that such canonical computations themselves could be embedded in canonical circuits^32,34^. The canonical circuit is thus portrayed as the structural component of the computational unit^33^ where slight modifications of either the hardware or software can shape a rich repertoire of network output^44,81^. Thus, an argument for straying away from the implementational level is rooted in computational identity of different circuits (across species) in the face of the multiplicity of their physical implementation^95,118^.

We suggest that for fine scale information (connectome or biophysically-detailed simulations) to provide any significant insight about the nature of computation, they should be further examined within a scale-dependent framework and considering:

> **The observation scale:** Although one can ignore detailed connectivity profile as unnecessary while considering the overall statistical properties as important, the proper choice of the scale remains crucial.
>
> **The structure-function relationship:** Physical details of the circuit affect the dynamics, suggesting that the structure constrains the computational function. However, constraining the space of possible computations does not provide an exact characterization of the performed computations because there exists no one-to-one mapping of structure-function.
>
> **Plasticity:** The ever-changing structure of the network implies a robustness in computational function despite changes in fine details of connectivity.
>
> **Subcellular control elements:** These elements (whether they are gap junctions, ion channels, dendritic spines, etc) provide mechanisms for change of function without apparent change of structure at the scale of microcircuitry.
>
> We have to recognize that since our system of interest harbors plasticity and homeostasis among other fundamental features outlined above (e.g. its hierarchy). These two aspects demand that our manipulations of the system be dynamic in nature in order to achieve that the intended effect is sustained over time despite the changing conditions (plasticity) and overall constraints (homeostasis).

## D. What does network control entail?

In the search for a proper spatiotemporal dynamics of control, some may resort to modern interpretations of network control. It is essential to recognize the short-comings of such an admixture of computational and implementational level. Although network structure determines certain properties of network dynamics such as limit-cycle oscillator synchrony^114^ or the likelihood of reliable dynamical attractors^51^, not all dynamics can be captured by the network structure alone. Some studies have considered a connectivity graph of complex networks equivalent to its dynamical nature, and have drawn the conclusion that recognizing the driver nodes is sufficient for understanding the strategy for controlling the network^68^. By reducing the dynamics to structural connectivity, they conclude that a large fraction of driver nodes (80%) are needed to control biological systems. Even though they point to the difficulty of controlling sparse inhomogeneous networks, in comparison to dense homogeneous ones^68^, their assumptions of reduction of dynamics to network structure and the definition of control and numerical methods in defining driver nodes are criticized by a) the evidence that a few inputs can reprogram biological networks^88^, b) trade-off between phase space nonlocality of the control trajectory and control input nonlocality^115^ and c) that the node dynamics – not degree distributions – define the nature of controllability^23^.

Another approach in advocating the structural control has been based on the assumption that active nodes can simultaneously activate all its connected neighbors but no further than that^89^. This assumption that is neither valid in the nervous system nor has any relevance to biological networks composed of elements with a variety of time constants and delayed communication. Additionally, not only the internal structure but also its connectivity pattern to the drivers and the depth of the network are essential in the emergent dynamics that follow stimulation. If the network is composed of a high external/internal node ratio, it will be largely influenced by the outside and its individual nodes will have a greater degree of independent behavior from the external stimulus. In contrast, if within system links are strong, then the system moves toward synchrony^21^. Likewise, the depth of the system is very crucial in its response to incoming stimuli. Systems with shallow depth are easier to force to behave in the desired way, as we can directly probe and influence the effects of external nodes on internal ones. A successful implementation of this principle is the first experimental evidence for the importance of individual neurons in *C. elegans* locomotion^126^. Using the wiring diagram of a shallow network (based on the only available full connectome, i.e. that of *C. elegans*^2,120^), it was predicted and confirmed that within a given class of motor neurons, only ablation of a subset should affect locomotion^126^. However, it was also noted that the connectome alone can not distinguish between different behaviors even with the same sets of input /output nodes. This issue is not surprising, since as we discussed in section 5, one of the characteristics of a given network (even) with fixed wiring, is its ability to manifest functional variability.

In contrast to shallow networks, deep networks are harder to control, and along each step we have to tweak the internal vs external nodes to achieve the desired outcome. One of the main practical challenges is that even controlling a “static” large network would require significant energy^92,115^. The requirement for high energy is due to the nonlocality of control trajectories in the phase space and its trade-off with the nonlocality of the control inputs in the network itself^115^. As a result, if the number of control nodes were to be constant, the required energy for driving the network scales exponentially with the number of network nodes^92^. This “required energy” exponentially decreases if the number of network nodes were to be constant while the number of driver nodes increases^125^. In addition, even if in certain cases network dynamics could be linearly approximated locally, two standing issues would remain that are not easily resolvable: first, control trajectories follow a non-linear mode and second, the local linear dynamics do not explain the major global properties of the network such as the basins of attraction^22,115^. Even if controllablity of static networks could be achieved, as soon as dynamic enters the game (in time varying systems), the required energy for controlling the system will make the control infeasble^125^. Overall, in high-dimensional systems that are governed by nonlinear dynamics, have dissipative properties (trajectories are confined only to a limited part of the permissible phase space), and where feedback imposes constraints on controllability, the mere identification of driver nodes and quantifying node variables are not sufficient to control the network^40,85^. Thus the control of complex networks requires both knowledge of the structure and dynamics across multiple scales of the system. To achieve multiscale control, and for not being obstructed by the inherent nonlinearity of the network dynamics, neuroengineers have to resort to designing system that are capable of providing compensatory perturbations in order to harness the nonlinear dynamics. This form of systematic compensatory perturbations is shown to be an effective tool in control of networks that manifest nonlinear dynamics^22^.

## E. Dynamical nature of computation and structure/function variability

The alterations in network behavior can originate from constant structural reshaping of the network (addition or deletion of links), or through modified intrinsic properties of the network nodes (i.e. neurons) and via changes in possible states of the interaction among the network nodes (i.e. synaptic weights). These different modes of reconfiguration of the network behavior set the dynamics of the information flow and the active behavior of the organism. A functional translation of these attributes leads to few rules, where a) one neuron can be involved in more than one behavior, b) one behavior can involve several circuits and c) one neuromodulator can alter multiple circuits. Naturally the involvement of these different modes in reconfiguration of the network behavior changes according to the size of the system and the repertoire of its functional states. In non-vertebrates with a very limited set of neurons, it is possible to pinpoint individual or several neurons as the key or even unique elements involved in highly reliable functions. For example, it is suggested that a single neuron in C elegans can be involved in highly reliable function^13^. In vertebrates (with larger brains), where routing and coordination of information flow becomes increasingly more important, local or global effects of neuromodulators and oscillatory rhythms play an essential role^52,61^. One can conclude that precision in control of the system depends on its complexity profile (i.e. the amount of information necessary to represent a system as a function of scale). As the number of neurons/circuits and diversity of network components increases, the reliability of functional dependence on individual components decreases. This increased complexity provides robustness at larger scale dynamics while it forgoes the details. Assessing the variability of structure/function thus becomes an essential aspect for directing stimulus at the right scale.

While the observed variability can be externally or intrinsically-driven^100^, in order to achieve functional adaptability in the light of external variability, the nervous system benefits from adapting stochastic responses^35^ where a given population activity can be construed as a likely sample from a posteriori distribution^48^. Intrinsic variability in the nervous system originates at different levels, from ion channels^37^ to synapses^31^, single cells^66,98^ and population level^12^. As a result, in driving the system, instead of aiming for a pre-determined response, the target should be to induce population activity such that it could be interpreted as one stochastic instance of the likely probability distribution. This supposed probability distribution of set of responses should depend on a multidimensional parameter space, where robustness and flexibility are attained through overlapping redundant functions. This form of redundancy is in stark contrast to engineered systems where multiple instances of identical copies are used to guarantee the robustness^72^. Instead, in the biological systems (and specifically in the nervous system), the robustness is achieved through a) variability at fine scales, where single cell variability exists in parallel with a robust representation at the population level^28,66,98^, b) networks with different configurations (of the underlying parameters) manifest robust response to neuromodulators^96^ and c) structurally different circuits respond reliably to external perturbation^71^. The existence of distinct stable basins of attraction despite the permissible extensive variability at fine scales, is an essential characteristic of complex systems. Such systems’ response to perturbation depends on the scale and the extent of the stimuli. In a sense, tolerance for small errors in complex systems comes at the price of the intolerance to large errors^60^. As a result, when the intrinsic variability is reduced due to excessively increased coupling, the abnormally greater ensemble correlation dramatically reduces the plethora of macroscopic states, a situation which is the hallmark of loss of complexity in organs with networks of excitable cells. Heart arrhythmia^93,94^ or seizures^28^ are examples of such loss of complexity.

## F. Are complex systems with fine-scale variability and robust macroscopic features predictable?

In simple chaotic systems, the lack of predictability is due to sensitivity to initial conditions. In contrast, in complex adaptive systems, information exchange across the scales is the main obstacle in the proper control of the system and the lack of predictability is due to relevant interactions and novel information created by these interactions. Ignoring the multiscale levels of information processing, one may jump to the conclusion that for proper control, we need to have a detailed knowledge of the individual elements, the initial conditions and their interaction among elements. Since we know that this is impractical if not impossible, can we do better? To properly control a system through stimulation, one has to have a proper knowledge of the interaction in the subunits of the system. However, only a controlled trajectory of the macroscopic states is desirable and yet achievable. From a dynamical system’s point of view, pushing the system at the right time and the right scale is the key to fine control. Given the scale-dependent interaction of the nervous system, we should ask: a) are complex systems with many scales inherently unpredictable? and b) if the answer is no, then how do we manipulate the system in a predictable way? Surprisingly, the answer to the first question is ‘’no, complex systems are not unpredictable”. However the predictability requires a few specific conditions and particulars. The existence of nonlinear feedback creates either a fast or slow ‘’wait-time of divergence” and therefore, up to some point, prediction holds and it rapidly deteriorates afterwards. This is similar to the behavior of some bifurcation systems. In fact, many systems that are considered to be periodic are inherently chaotic. A prime example is the solar system where its very slow divergence time (4 million years) leads to the observation of chaotic behavior only when 100 million years of the entire solar system are examined^116^. Therefore to navigate the system in the aimed trajectory, we have to constantly adjust and push the system such that it does not deviate from the desired path after the implementation of the last stimulus. This strategy can be effective only if the system does not have low-dimensional chaotic behavior. Otherwise, any simple perturbation could lead to unrecoverable massive perturbation. In the case of the ensemble activity (such as cortical processing with the involvement of a much larger set of neurons in comparison to the simple nervous system of invertebrates such as C Elegans), this notion becomes highly relevant since low dimensional neural trajectories provide surprisingly accurate portrait of the circuit dynamics^98^ and dimensionality reduction methods exhibit effectiveness in population decoding^24^.

In non-chaotic system, our ability to modify the system is limited^106^. If the system’s behavior is periodic or multiply periodic (nested periodicity), we are stuck with the system’s intrinsic periodicity dynamics. We can slightly change the orbit by small perturbations or we can only induce perturbations that would switch the dominant periodicity from one of the existent orbits to another existent one. This limitation also means that the achieved effect is robust because of the existent orbits, but it will not permit large alterations in the system. In contrast, the existence of chaos could be helpful in controlling a chaotic system. Since systems with a chaotic attractor have an infinite number of unstable periodic orbits, one can exploit this property and push the system towards one of the already existing unstable periodic orbits^91^. However, if the system is high dimensional evolving chaotically on a low dimensional attractor, this method no longer works. The reason for the ineffectiveness of this method in this case is that one would require an excessive amount of information extracted from the data and a very long history of the dynamics in order to be able to properly achieve control^4^. Though, interestingly, the situation is not hopeless, as one can exploit the low-dimensionality of the system dynamics and through a feedback control and repeated application of tiny perturbations, control the system^4^. Here we advance the idea that recurrent feedback in the neuronal networks might indeed be the evolutionary mechanism developed to specifically deal with the low dimensional dynamics of a high dimensional system for providing intrinsic control for response reliability. This method has been extended to control excitable biological systems in the past. It has been shown that by irregular (based on the chaotic time) delivery of the electrical stimulation, the cardiac arrhythmia can be pushed back to a low-order periodic regime^39^. In the hippocampal slice (CA3), the same method of entrained spontaneous burst discharges proved to be much more effective than periodic control^103^. Other methods, such as periodic^5,63^ or stochastic^36^ stimulation can affect chaotic systems too but it is hard to predict their effect for networks with many layers.

## G. Precision and reliability in control of neuronal networks

Perturbation due to noise and presence of variability are the key challenging issues in the control of chaotic (recurrent) neural networks. Recurrent networks operating at high gain, even without any stimulation, show complex patterns and chaotic dynamics that are very sensitive to noise^113^. In high-gain regime, small changes in synaptic strength of recurrent networks can easily lead to chaotic instability, lack of robust output and high sensitivity to noise^6,113,121^. This behavior is in contradiction to biological networks which show robustness in the presence of high synaptic plasticity^20^. As a result, while recurrent networks are potentially computationally powerful systems, their unreliable behavior, in the face of noise and synaptic plasticity, renders them ineffective models of their true biological counterparts. Despite these challenges, some solutions have been proposed to tame the chaos in firing rate recurrent networks. By tuning the weight of the recurrent connections, i.e. minimizing the error between a desired and the current trajectory, robustness can be achieved^67^. This tuning leads to a regime that exhibits chaotic dynamics and locally stable nonperiodic trajectories, allowing reproducible behavior^58^. The stable trajectories act as dynamic attractors enabling the network to resist deviations in response to perturbation. Unfortunately, this well-desired property of robustness is very hard to achieve in spiking recurrent networks where the irregular firing states manifest intense chaos. Both experimental (rat barrel cortex under anesthesia)^69^ and theoretical^82,83^ studies show that a single spike can rapidly decorrelate the microstate of the network. Surprisingly, state perturbations decay very quickly and single-spike perturbations only lead to minor changes in the population firing rates^83^. Yet still, these complex but unstable trajectories always lead to exponential state separation and, as a result, these network can not reliably map the same input to nearby neighborhood in population trajectory. In addition, *in vivo* non-anesthesized neurons have higher conductance^30^ in comparison to the anesthesized state, where the spike-induced stability has been observed experimentally^69^. Therefore, it is highly likely that the perturbation induced instability is much more intense in the presence of higher intrinsic noise in non-anesthesized cortex.

In the face of complex network topology and synaptic plasticity, reliability of continuous nonlinear networks and their controlled perturbation remains an unsolved challenge. Sensitivity to the input, insensitivity to perturbations and robustness despite changes in initial conditions are the requirement of stable yet useful computation^58^. These attributes are much easier to achieve in the idealized classical model of neural state space^47^, where the stability of computation arises due to convergence to steady-state patterns as fixed-point attractors. Yet higher cognition requires “efficient computation”, “time delay feedback”, the capacity to “retain information” and “contextual” computation^29^. These desired properties are more in tune with the characteristics of reservoir computing (rather than static attractors), namely “separation property”, “approximation property” and “fading memory”^29^. However, the transient dynamics of such liquid-state-like machines^70^ require a sequence of successive metastable (“saddle”) states, i.e. stable heteroclinic channels^98^ to manifest robust computation in the presence of noise and perturbation. The necessary condition for transient stability in a high-dimensional systems with asymmetric connections is to form a heteroclinic sequence linking saddle points^98,99^. The successive metastable states ensure that all the trajectories in the neighborhood of these saddle points tend to keep the flow of computation in the desired channel. This property can be harnessed to ensure robustness of controllability despite the inherent variability of transient dynamics. To achieve proper control, instead of resorting to manipulation of the target behavior at its end-point, it is more efficient to apply minimal, yet repeated stimuli that can minimize the deviation of trajectories between the metastable states. While it remains impossible to control continuous nonlinear networks, targeted control at multiple scales seems to be a good remedy and solution where neuroengineers should focus.

The limitations for proper control of abstract and simple neural network models may lead to the conclusion that infinite precision is required to fully describe or control the computation of analog neuronal networks with unbounded precision. Given that infinite precision would turn to nontrivial nonlinearity, one could infer that the proper control of natural neuronal networks would be hard/impossible to achieve. Yet the nervous system mysteriously manifests routine robust macroscopic behavior. This puzzling dilemma hints on some possible solutions. Surprisingly, it has been shown that linear precision is sufficient to describe (up to some limits of) analog computation^9,109^. This linearity would map to N bits of weights and N bits of activation values of driver neurons for up to the Nth step of the computation. Following this principle, “analog shift map”, a dynamical system with unbounded (analog) precision was designed and shown to portray computational equivalence to the recurrent neural network; yet its tuning only required linear precision^108^. Therefore it seems likely that the nervous system reaches reliability by bounding its analog precision to modular computation, i.e. computation at scale. This multiscale property of neural computation means that internal control is achieved by decomposition of tasks with many needed cycles of computation to smaller modules in order to keep the tuning linearity in place. Equivalently, to achieve control during perturbation, one needs to pay attention to the computational limits at each scale, where no computational procedure of N steps could be mapped to more than N bits of weights and N bits of activation. Any attempt to control the system at higher precision will not change the result and lesser precision will fail to achieve control. Perhaps this computational principle is also among the reasons that the primitive nervous systems are mostly formed as diffuse nerve nets, while more advanced nervous systems harbor a multi-scale structure to fully benefit contextdependent information processing^27^. In engineering neural control systems, one has to pay close attention to the achievable precision at each scale, its indifference to small values, yet the saturation as measured as a function of the computation time within that scale.

## H. Concluding remarks

In linking the structure and dynamics of the neuronal network, mapping the space of possible interactions becomes important and a matter of challenge. Since all the possible interactions at a fine scale occupy a vast multidimensional space, in order to have robust behavior, the system is likely to rely on a more limited set of probable interactions between local densities (i.e. ensemble activity). As the scale grows, the set of possible interactions decreases yet the outcome of such interactions better matches with the behavioral (macroscopic) outputs of the system. Relevant parameters are those permissible sets of interactions that have increased probability of occurrence as the scale increases. Functional state transitions depend on the spatial variation and interaction of the ensemble activity densities. As a result, the complexity profile defined as the amount of information necessary to represent a system as a function of scale gives us the number of possible states of the system at a particular scale^7^. Therefore the finer the computational scale of the system, the more information is needed to describe it. It is through these mechanisms that the system can maintain its intrinsic dynamical balance yet manifest responsiveness across multiple time scales^28^ and provide stereotypical macroscopic spatiotemporal patterns in the lights of microscopic variability^61^. This viewpoint defies the blind big-data approach of incorporating more details with the hope of yielding a better model and more accurate prediction^8^. Even with the assumption that at some point in the future we can map the connectivity and the activity of all the elements of the neuronal network, we are still in need of an information-theoretic formalism that shows how the output is sensitive to fine scale perturbations, and how the coarse-scale reflects redundancy and synergy of the aggregate activity of finer scales^25^. In order to enhance the effectiveness of targeting multiscale neural systems, alterations of biophysically-based features should match the level of the desired computation. In targeting a stream of computation, the link between the computation and the architecture is what defines the optimal solution for maximizing the efficiency of the stimulation. In summary, to better control the system, we have to focus on the information transfer across multiple scales. Only with this approach can engineering advancements in precise opto-electric stimulation open ways to alter the system in the desired way.

